# Sub-clinical glutamate receptor antagonist combinations prevent progressive demyelination

**DOI:** 10.1101/2024.03.19.585708

**Authors:** Verity F. Mitchener, Chris Bulman, Alexander Mellor, Nishita Bhatt, Badrah Saeed Alghamdi, Jackie Nash, Rhianon Proctor, Nancy Smith, Abbie Newman, Victoria Nwabiakam, Gideon Stone, Daniel Morgan, Waldemar Woznica, Glenn Harper, Mario Valentino, Alberto Pérez-Samartín, Fernando Perez-Cerdá, Carlos Matute, Robert Fern

**Author notes:** **Materials requests and Correspondence** should be addressed to Robert Fern, Peninsula Medicine School, University of Plymouth, John Bull Building, Research Way, Plymouth, PL6 8BU, UK.

## Abstract

Multiple sclerosis (MS) affects almost 3 million people globally who suffer demyelination as a series of relapses and remissions that tend towards progressive deterioration over time. The proximate cause is auto-immune attack by the adaptive immune system; therapies directed against this are effective during the relapsing-remitting phase but are less effective or ineffective during progression where other injury mechanisms may be significant. LPS- and cuprizone-induced experimental demyelination share features of progressive demyelination in MS but the underlying mechanisms are not well understood. We show here that these demyelination models can be reproduced *ex vivo* using short protocols, revealing that combined antagonism of two types of glutamate receptor, NMDA and AMPA, using clinically approved antagonists at sub-clinical doses, can protect against these forms of demyelination. Combined low dose therapy was subsequently shown to be effective *in vivo* against dietary cuprizone and experimental autoimmune-encephalomyelitis (EAE) models of demyelination and, in particular, protected the smaller myelinated axons that are the main substrate of the function loss in progressive MS.

---

MS is a chronic demyelinating disorder that progresses via complex and only partially understood mechanisms. The initial pathogenic trigger involves targeting of central myelin by the adaptive immune system, but aberrant innate immune responses and glutamate receptor over-activation are also implicated ^1–4^. These two components may be most significant in the progressive phase of the disease which is refractory to established therapies directed at components of the adaptive immune system.

The main tools in the study of MS are rodent EAE models that recapitulate many of the clinical, cellular, and structural features. An alternative model of demyelination and subsequent remyelination involves dietary supplementation with the toxin cuprizone (CPZ), which acts via a largely unknown mechanism. These animal models involve long *in vivo* protocols and are useful primarily for low-throughput studies. There is a compelling need for higher throughput assays to expand both the number of drugs tested and the range of doses examined. EAE models recruit an adaptive immune response against central myelin to produce a gradual functional loss that becomes apparent 1-2 weeks after inoculation. In contrast, the myelin damage associated with aberrant innate immune responses or CPZ diet develop more rapidly. Innate immune reactions are initiated by Pathogen-Associated Molecular Patterns (PAMPs) such as the bacterial toxin lipopolysaccharide (LPS). Systemic LPS inoculation produces functional disruption of adult white matter within 3 days ^5^, and direct LPS injection into adult spinal cord produces focal demyelination over the same time-course ^6^. In the standard CPZ diet, molecular disruption of myelin and oligodendrocyte apoptosis is evident from day 2 ^7–9^, and myelin vacuolization is significant after 1 week ^10^.

## New *ex vivo* models of acute demyelination

Myelin damage is an early feature of both MS and its EAE models and is apparent before any structural change to oligodendrocyte somata ^11^. Under *ex vivo* conditions where penetration through the blood brain barrier is not an issue, we found that 60 min exposure to 1 μg/ml LPS produced a fall in compound action potential (CAP) conduction in CD-1 mouse optic nerve (MON) to 75.2 +/-5.1% vs. control. The CAP continued to decline over a subsequent 60 min period of LPS washout, reaching 56.1 +/-6.3% (P<0.001 vs control; Fig 1 a, red data). No significant difference in the rate or extent of action potential loss was produced by a 120 min LPS exposure (black data), showing that 60 min of LPS was sufficient to recruit a mechanism that led to progressive loss of function. This protocol produced myelin pathology including low morphology scores and lamella splitting and decompaction that was apparent as an increase in myelin sheath thickness and periodicity, with a corresponding reduction in axon g-ratio with no change in axon diameter or peri-axonal space area (Fig 1 b, c, Ext. data Fig 1). Using a novel hemi-sected brain slice protocol (see Methods) applied to PLP-GFP mice, 60 min 1 μg/ml LPS + 60 min recovery reduced the fluorescent signal from the vital myelin dye Fluoromyelin Red (FM-Red) in the corpus callosum (MCC) by 46.5 +/-3.6% (Fig 1 d), indicating molecular disruption of myelin ^7^. In the same sections, LPS had no lytic effect upon GFP(+) oligodendrocyte somata, but produced disintegration and reduced fluorescence in GFP(+) cell processes (Fig 1 d). This acute LPS protocol also generated a robust reactive astrocytosis (Ext. data Fig 2 a, b), and the loss of oligodendrocyte processes was confirmed in the absence of the FM-red staining step (Ext. data Fig 2 c, d). Immuno-staining against LPS in tissue fixed following the LPS protocol showed no or low levels of binding was observed in PLP-GFP expressing oligodendrocytes, some LPS binding to GFAP(+) astrocytes and extensive binding to Iba1(+) microglial cells (Fig 1 e). This corresponded with a significant shift from Iba1(+) microglia with a ramified morphology typical of the resting state, to morphologies typical of bushy, motile, and phagocytic cell forms (Fig 1 f). LPS is a large molecule and penetration into the tissue and cellular binding was confirmed in PLP-GFP(+) mice using a fluorescently tagged analogue (Fig 1 g), and using antibody staining of untagged LPS in sections unexposed to other antibodies (Ext. data Fig 2 e).

**Figure 1.**
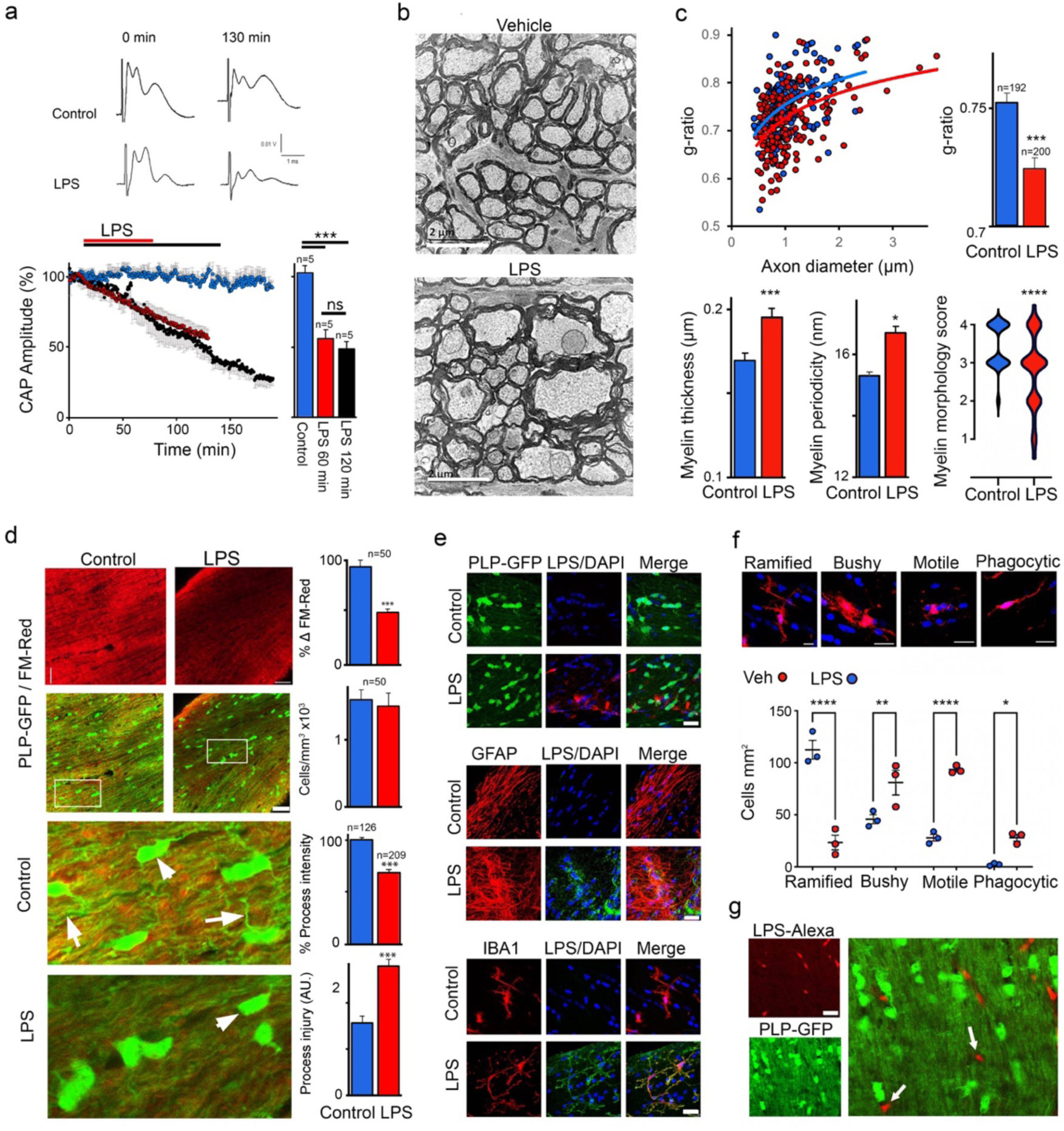
The innate immunity activator LPS rapidly damaged central myelin. a: A standard LPS protocol (60 min 1 mg/ml LPS + 60 min wash-out; red data and bar), produced a 56.1 +/-6.3% decline in the action potential (top: representative CAPs). Extending the LPS exposure to 120 min had no significant additional effect at this time point. b, c: The CAP loss was associated with myelin decompaction evident in representative x-s ultra-micrographs (b), and in a reduction across the axon diameter spectrum in g-ratio (c, top), and in an increase in mean myelin sheath thickness, myelin lamella periodicity, and myelin injury score (c, bottom). d: Left: the standard LPS protocol produced a significant loss of signal from the fluorescence myelin dye FM-Red (top), and of GFP from processes of PLP-GFP oligodendroglia (boxed areas shown under at higher gain), which developed a damaged appearance. Right: Data summary (n=ROI). e: Following wash-out, immuno-detection of LPS binding showed no or low levels of colocalization with GFP(+) oligodendroglia (top), a degree of colocalization with the astrocyte marker GFAP (middle), and extensive colocalization in Iba-1(+) microglial cells (bottom). f: Top: The four Iba-1(+) microglial morphology types found in white matter (red). Bottom: The proportion of ramified microglia significantly declined following the LPS protocol, and bushy, motile and phagocytic forms significantly increased. g: Alexa-568-conjugated LPS (red) bound to numerous GFP(-) glial somata in PLP-GFP MON. Scale bar = 2 μm (b), 10 μm (d-g).

**Figure 2.**
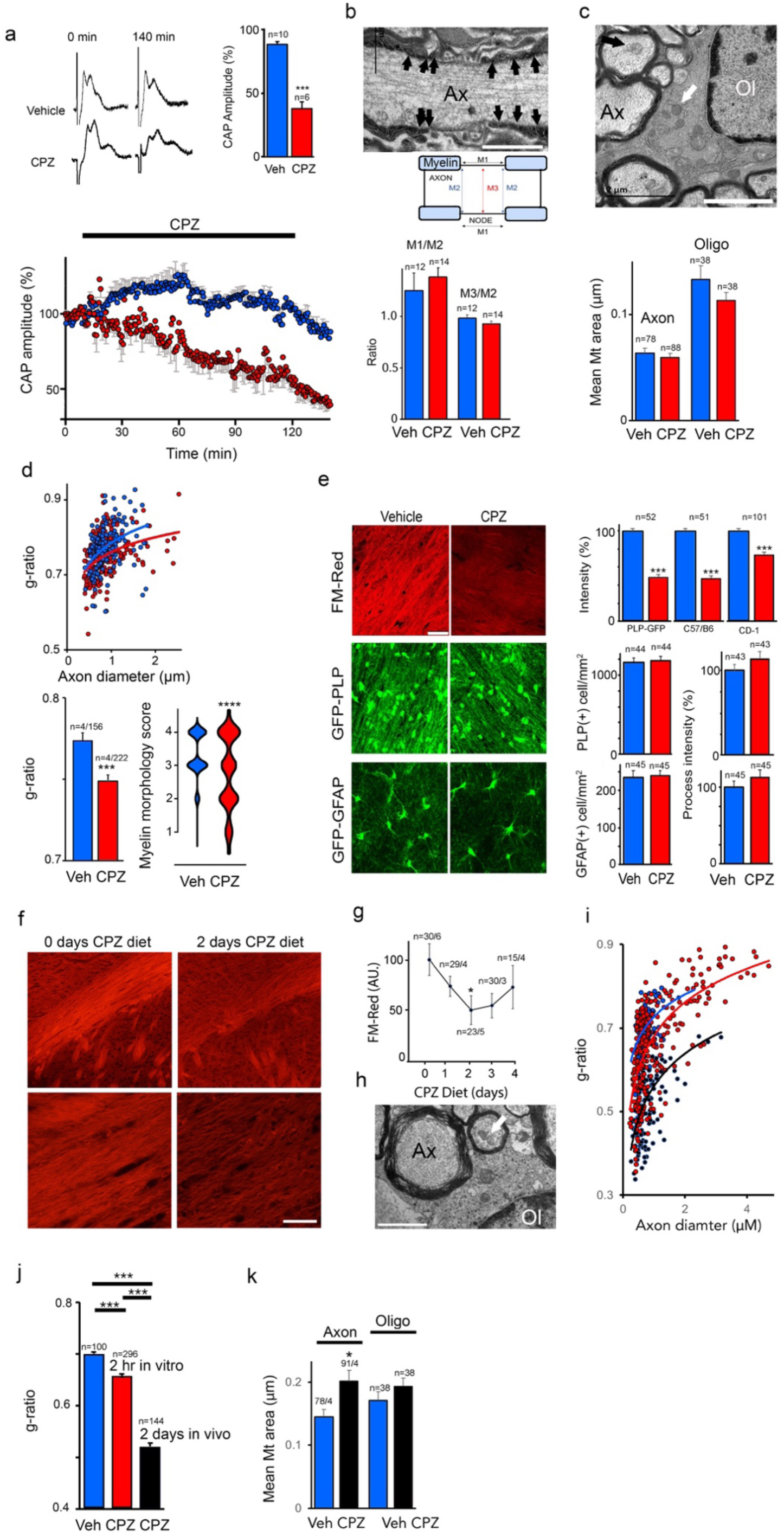
CPZ rapidly damaged central myelin. a: 120 min perfusion with CPZ (black bar) produced a significant loss of the CAP (to 52.07 +/-0.07% after 30 min recovery, blue data points). b (top): a typical node of Ranvier (black arrows = paranodal end loops; insert shows the morphological features analysed; scale = 1 μm). b (bottom): CPZ treatment had no significant effect on node morphology. c (top): Typical Oligodendrocyte (‘Ol’; white arrow = mitochondria), surrounded by axon profiles (‘Ax’; black arrow = mitochondria). b, c, bottom: CPZ treatment had no effect on mitochondrial area nor the morphology of the node. d: CPZ reduced axon g-ratio and myelin injury score. e: CPZ significantly reduced the fluorescent FM-Red signal in three mouse lines, but had no effect upon the number of GFP(+) somata or processes in either PLP-GFP or GFAP-GFP mice. f: 2 days of CPZ diet in C57BL/6 mice produced a significant loss of FM-Red signal. g: Time series data for different lengths of CPZ diet. h: 2 days of CPZ diet produced myelin decompaction in axons, and no oligodendrocyte pathology in the optic nerve (white arrow = oligodendrocyte mitochondria, scale = 1 μm). i, j: Comparison of the g-ratio of axons in the optic nerve of C57BL/6 mouse subject to the 60 min isolated CPZ (red) or vehicle (blue) protocol, or to 2 days of CPZ diet (black). k: Mitochondria were significantly larger in MON axons following 2 days of CPZ diet, but not in oligodendrocytes. Scale bar = 1 μm (b, c, h), 10 μm (f).

Acute exposure to CPZ *ex vivo* had a similar effect to LPS, producing a decline in action potential conduction to 38.6 +/-5.2% after 120 min plus 30 min reperfusion (P<0.001 vs vehicle; Fig 2 a). The standard CPZ diet model produces mitochondrial disruption ^12^, but the rapid functional loss we report here *ex vivo* was not associated with any significant disruption in the structure of the node of Ranvier (Fig 2 b), which is an early sign of metabolic disturbance in white matter, or significant mitochondrial swelling in either axons or oligodendroglia (Fig 2 c). As with LPS, the functional disruption produced by CPZ was associated with decompaction of axonal myelin that was evident as a decline in the fibre g-ratio and myelin morphology score (Fig 2 d). No effect was produced by perfusion with the 0.1% DMSO vehicle (Ext. data Fig 3). CPZ exposure reduced the myelin FM-Red signal to 74.3 +/-2.5% of vehicle control in the CD-1 mice used for all the experiments on non-transgenic mice to this point, compared to vehicle controls. In PLP-GFP transgenic mice the reduction was to 50.6 +/- 2.5% and was 48.8 +/-1.9% in C57BL/6 mice that are the wild-type background for these transgenic mice (Fig 2 e). In PLP-GFP mice, CPZ had no significant effect upon GFP(+) oligodendroglia somata, or of GFP(+) astrocyte somata in GFAP-GFP mice (Fig 2 e). This protocol also had no significant effect on the ability of either type of glial cell processes to retain GFP (Fig 2 e; Ext. data Fig 4), consistent with their undamaged ultrastructural appearance. In a C57BL/6 mouse CPZ diet model, we found significant loss of FM-Red emission in optic nerve fixed after 2 days of diet (to 61.2 +/-6.0%, Fig 2 f, g). Similar myelin decompaction was found after the standard *ex vivo* model in the C57BL/6 mouse and after 2 days of CPZ diet (Fig 2 h-j), but the effect was more pronounced in the diet group where g-ratio declined to 0.52 +/-0.004. No significant mitochondrial swelling was found in the oligodendrocytes of CPZ diet-fed mice, but an increase in axonal mitochondrial size was seen that may indicate early metabolic stress (Fig 2 h, k).

**Figure 3.**
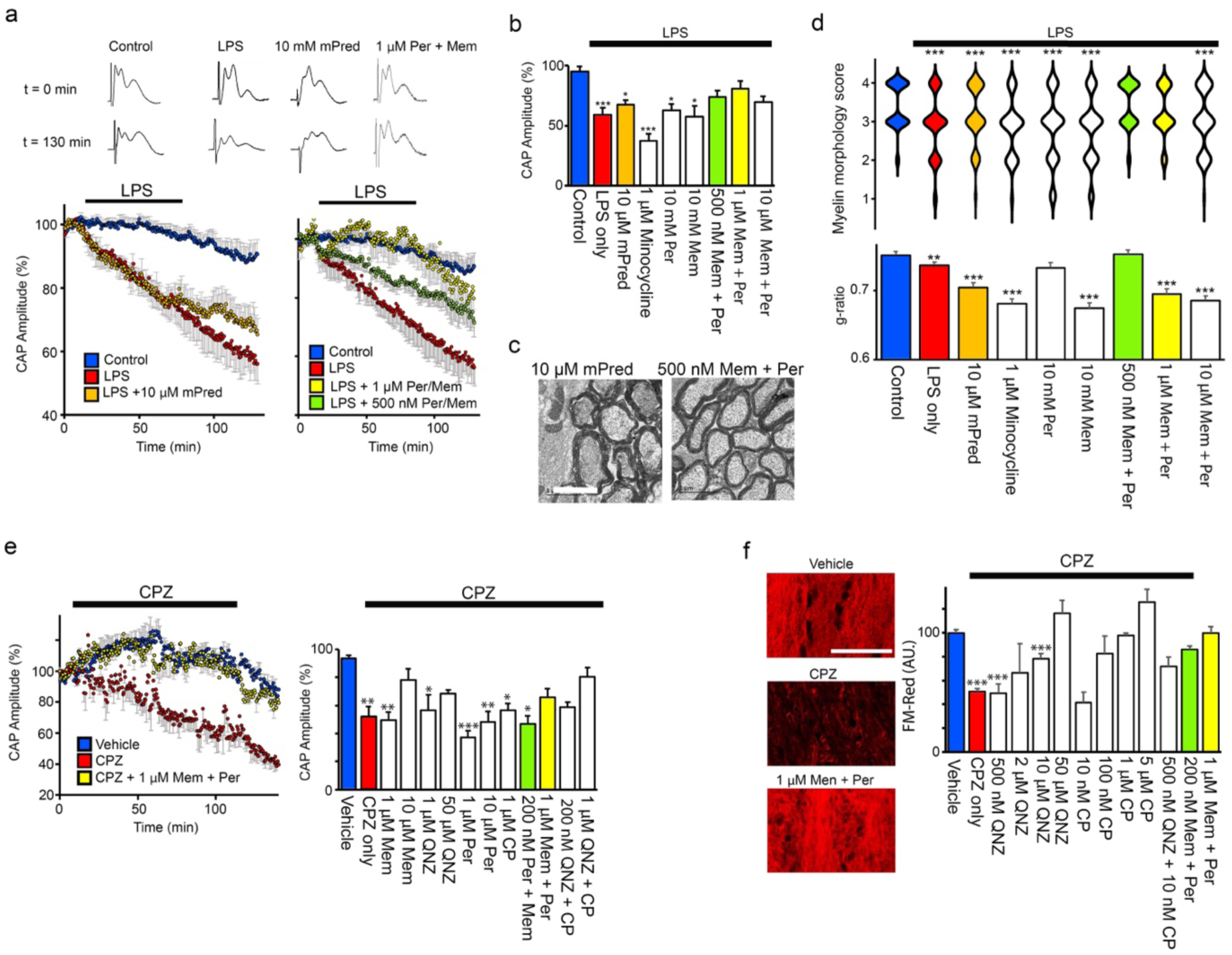
LPS and CPZ act via a common excitotoxic mechanism. a: The disruption of action potential conduction produced by LPS is not affected by methyl prednisolone (mPred), but is largely prevented by a combined low-doses of clinically-approved glutamate antagonist (CLD CAGA) protocols based on memantine and perampanel (top: representative CAPs, Bottom: mean CAP amplitude data). b: CAP data summary. c: Myelin decompaction in LPS-treated MON axons is evident in the presence of mPred but not in CLD CAGA conditions. d: Blind myelin injury score distribution (top) and axon g-ratio (bottom) changes evoked by LPS are normalized by CLD CAGA protocols but not by anti-inflammatory compounds mPred and minocycline. e: CPZ-induced action potential disruption is largely prevented by CLD CAGA, and by a similar paradigm using the experimental glutamate antagonists QNZ-46 and CP465022. f: The FM-Red fluorescence loss evoked by CPZ is prevented by the selective NMDA and AMPA antagonists QNZ-46 and CP 465022 at high concentrations, and by low combined treatments including CLD CAGA (Mem + Per). Scale c = 1 μm; f = 10 μm.

**Figure 4.**
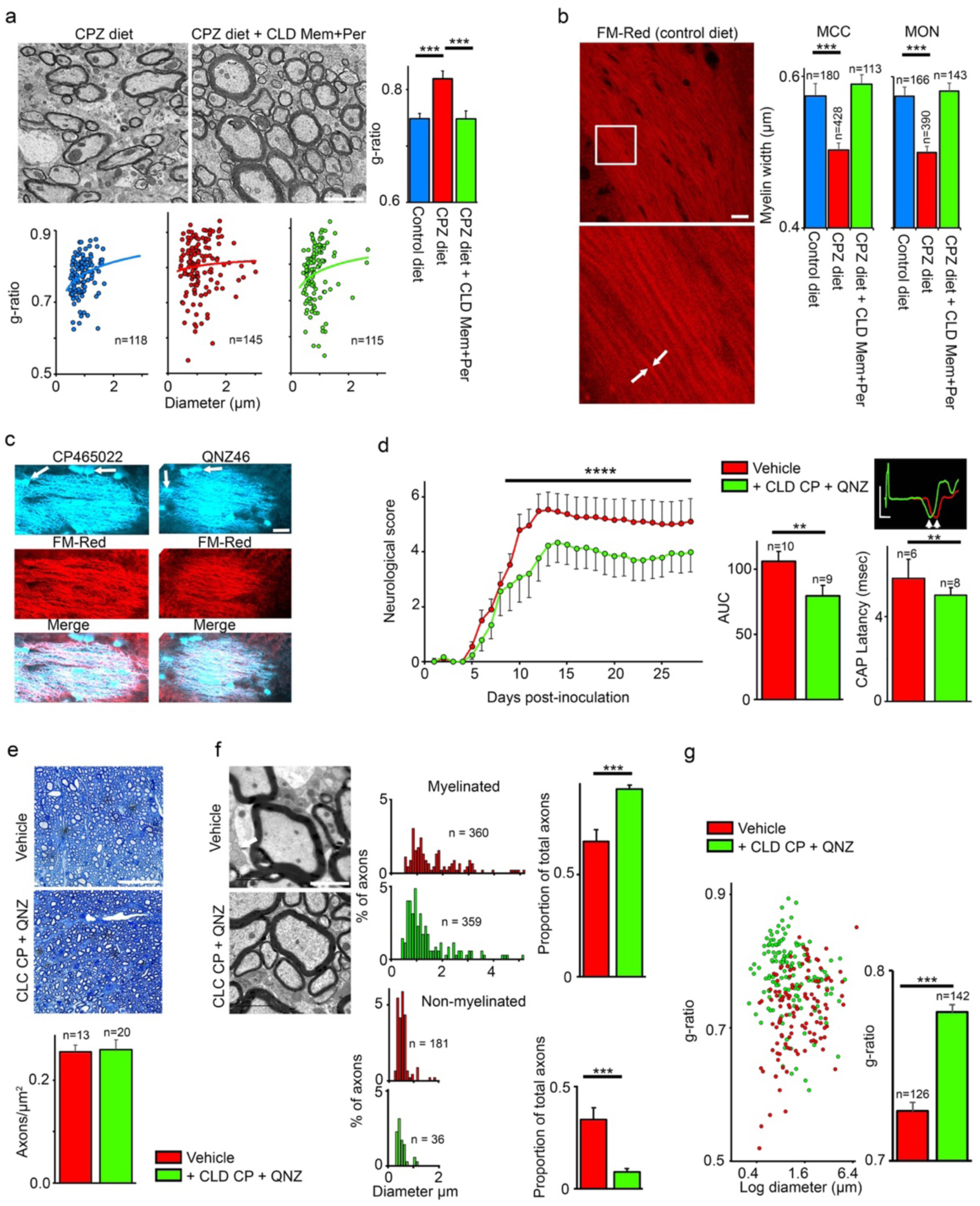
CLD-GA treatments are effective in *in vivo* models of demyelination. a: Daily Mem + Per CLD treatment reverses the increase in MCC axon g-ratio produced after 28 days of CPZ diet (scale = 2 μm). b: This treatment also reversed the corresponding thinning of myelin sheaths produced by the diet in MCC and MON, assessed via FM-Red confocal imaging of individual sheaths (eg., white arrows; scale = 10 μm, boxed area under). c: QNZ46 and CP465022 fluorescence following UV excitation in MON following CLD-GA treatment based on these antagonists (top), and the corresponding FM-Red co-staining (middle), showing co-localization (scale = 10 μm). Note oligodendrocyte somata with processes contiguous with myelin sheaths also concentrate the drugs (white arrows). d: CLD-GA daily treatment with QNZ46 + CP465022 significantly reduced the clinical severity in EAE (left: time-course, Mann-Witney U-test; right: Area under the curve, T-test), and improved the conduction velocity of spinal cord axons (CAP peaks indicated by white arrows, scale = 1 mV/msec). e: CLD CP 465022 + QNZ46 did not affect spinal cord axon density in EAE (n= section, scale = 20 μm). f: This treatment eliminated a large population of small naked axons in EAE spinal cord (left, white arrows), and proportionally increased the population of corresponding small myelinated axons (scale = 1 μm). g: The g-ratio of spinal cord axons in treated mice was higher in smaller diameter axons (note log scale).

## A novel AMPA/NMDA receptor parallel injury pathway

LPS is known to rapidly recruit pro-inflammatory cytokine / nitric oxide release pathways in both microglia and astrocytes that are potentially damaging to white matter ^13–15^. These events are sensitive to the anti-inflammatory drugs minocycline and methylprednisolone ^16^, but these drugs had no significant effect on the functional and structural pathology evoked by 60 min LPS exposure (Fig 3 a-d; Ext. data Fig 5). In contrast, combined low dose-glutamate antagonist (CLD-GA) protocols based on clinically approved drugs (500 nM or 1 μM of the AMPA receptor antagonist perampanel + 500 nM or 1 μM of the NMDA receptor antagonist memantine) protected against LPS-induced action potential loss (Fig 3 a, b; Ext. data Fig 6), myelin decompaction, and myelin injury (Fig 3 c, d).

Mono-treatment with 10 μM perampanel was protective against decompaction, but not against myelin injury or action potential loss, and 10 μM memantine alone had no significant protective effect (Fig 3 b, d; Ext. data Fig 7). Prior studies have shown that 10 min of LPS exposure is sufficient to elevate extracellular glutamate in isolated CNS preparations via poorly understood but generally cytokine-independent mechanisms ^17–19^. CPZ has a similar effect upon extracellular glutamate ^12^ and we found that CLD-GA protocols were protective against the functional loss (Fig 3 e) and myelin disruption (Fig 3 f) produced by CPZ ex vivo. For example, 1 μM perampanel + 1 μM memantine significantly reduced the action potential decline (Fig 3 e, right), and eliminated the disruption to myelin structure assessed via FM-Red imaging (Fig 3 f). The experimental glutamate antagonists QNZ-46 and CP465022 were also highly effective at providing functional protection at low combined doses (Fig 3 e).

## Combined low dose glutamate antagonist therapy

We tested the CLD-GA approach in established *in vivo* models of demyelination. We employed a 28-day CPZ diet paradigm where myelin injury in the MCC and MON is extensive but re-myelination is limited ^20,21^. Myelin pathology was quantitated as an increase in axon g-ratio in the MCC (Fig 4 a), and as thin myelin sheaths assessed via confocal FM-Red imaging in MCC and MON (Fig 4 b), two well-characterised features of myelin pathology in this model. A CLD-GA treatment regime based on sub-clinical doses of clinically approved therapies (2 mg/kg/day memantine + 0.8 mg/kg/day perampanel) effectively prevented the myelin damage (Fig 4 a, b). CPZ is a copper chelator, but the demyelinating effect is not mitigated by copper administration ^9,22^. Morphological changes to mitochondria have been reported, but isolated mitochondria are not affected by the toxin ^9^, and the mode of action of the toxin is therefore not well established. Recent studies have provided convincing evidence for a role for ferroptosis ^8^, a form of cell death triggered by iron influx where free radicals produce lipid peroxidation. Ferroptosis is an important down-stream pathology associated with glutamate excitotoxicity ^23,24^, and the current findings identify a form of glutamate excitotoxicity mediated by parallel activation of AMPA and NMDA receptors as the mechanisms underlying CPZ toxicity both *ex vivo* and *in vivo*.

Extracellular glutamate is elevated in both normal-appearing white matter and lesion sites in MS patients ^25,26^, and the concentration of the neurotransmitter rises at lesion sites 6-12 months before the lesions themselves appear ^27^. Monotherapy with NMDA or AMPA antagonists is protective in EAE models of MS, but in general only at high concentrations ^3,28–31^. For example, the protection afforded by memantine requires a dose of 60 mg/kg/day, while a dose of 10-20 mg/kg/day in rodents correlates with the clinical guidelines for memantine in humans ^32^, a dose that has no benefit in clinical trials of MS patients ^33^. The memantine/perampanel CLD-GA paradigm that is effective against CPZ-induced white matter injury was based on doses an order of magnitude below the clinical treatment range for these drugs and is an attractive alternative to full-dose co-administration such as that tested in stage II clinical trials for glioblastoma (eg., ^34^). Membrane-to-channel inhibition has recently been described for lipophilic antagonists of NMDA and AMPA receptors, which can diffuse from the cell membrane into the receptor protein via gated fenestration ^35,36^. To take advantage of this phenomena and further extend the clinical potential of the CLD approach, we tested a combination of two experimental antagonists that are structural analogues of myelin-targeting fluorescent dyes.

QNZ46 and CP465022 are fluorescent 3(4H) quinazolinones that incorporate the myelin-targeting trans-stilbene pharmacore ^37^. QNZ46 is a selective blocker of NMDA receptors incorporating the GluR2C/D subunit such as those found in myelin ^38–40^, and CP465022 is selective for AMPA receptors ^41^. These lipophilic drugs fluoresce blue upon UV excitation allowing their localization at the sub-cellular level ^42^, and revealing their selective uptake into myelin and oligodendrocyte soma in white matter (Fig 4 c). No such fluorescence was detected in cellular components that were not contiguous with FM-Red stained myelin and these drugs therefore offer myelinating cell targeting and were tested at 2 mg/kg/day QNZ46 + 1 mg/kg/day CP465022 in an EAE model. This combined dose is an order of magnitude below the established *in vivo* mono dose of each ^31,42^, and significantly improved both the neurological score and the evoked action potential latency in EAE compared with vehicle treated mice (Fig 4 d). Demyelinated and non-myelinated axons cannot be reliably distinguished in cross-section ultra-micrographs, but prior studies report that non-myelinated axons make up ∼10% of spinal cord dorsal column axons (eg., ^43^ ^44^). In EAE mice, 33.9 +/-5.9% of axon profiles in the spinal cord dorsal column were un-myelinated in the vehicle group, and this fell to 8.2 +/-1.7% in the drug treated animals (Fig 4 e, f). The corresponding increase in the proportion of myelinated axons and the significantly higher g-ratio in the drug-treated mice, in particular of smaller diameter axons (Fig 4 g), confirms the extensive protection of the small-diameter myelinated axons that are known to be selectively damaged in progressive MS ^45,46^.

Myelinic NMDA receptors sense activity-dependent glutamate release from axons and regulate the delivery of energy substrate from the sheath to the axon cylinder ^47^. Myelin also contains AMPA receptors that similar to NMDA receptors ^38–40^ can act as a significant route of damaging Ca^2+^ influx when over-activated ^48,49^. Unlike synaptic receptors, myelinic NMDA receptors are only weakly sensitive to Mg^2+^-block ^39,50^ and are therefore unlikely to require collateral depolarization by non-NMDA glutamate receptors for functional activation ^51^. Even small rises in glutamate concentration such as those reported in pre-lesion white matter of MS patients (∼1 mM, ^27^) may therefore activate NMDA and AMPA Ca^2+^-influx pathways and trigger the novel parallel injury mechanism we have described in central myelin. The current findings show that partial block of both receptors significantly reduced myelin injury. Prior studies using EAE models report a protective effect of AMPA and NMDA antagonist monotherapy. These studies have not translated clinically, most plausibly because the high antagonist concentrations required for monotherapy protection are not suitable for the management of a chronic human disease. The development of *ex vivo* versions of two alternative models of demyelination has allowed us to identify a new therapeutic approach with no predicted side-effects suitable to prevent acute relapsing demyelinating episodes that are intractable to established therapies.

## Methods

### Animals and reagents

All animal procedures conformed to local ethical standards and ARRIVE guidelines. UK home office and Spanish national regulations were followed as appropriate. The majority of mice used in the experiments were female, but mixed populations of transgenic animals were employed. For isolated MON and MCC experiments, brains were collected from adult (P80–110) CD-1, heterozygous PLP-GFP, or C57BL/6 mice. For MCC, 250µm vibratome-cut coronal sections were cut in oxygenated, ice-cold cutting solution (in mM): NaCl, 92; KCl, 2.5; NaH_2_PO_4_, 1.2; MgSO_4_, 2; CaCl_2_, 2; NaHCO_3_, 30; glucose, 25; Hepes, 20; Na Pyruvate, 3; Thiourea, 2; Na Ascorbate, 5; pH, 7.4. Acute experiments using MON and MCC were performed in artificial cerebrospinal fluid (aCSF), composition (in mM): NaCl, 126; KCl, 3; NaH_2_PO_4_, 2; MgSO_4_, 2; CaCl_2_, 2; NaHCO_3_, 26; glucose, 10; pH, 7.45, bubbled with 5% CO_2_/95% O_2_ and kept at 37 °C. Lipopolysaccharide (LPS) from *Escherichia coli* was purchased from Sigma Aldrich (O111:B4, Batch #019M4009V) (UK), cuprizone (CPZ) from Sigma Aldrich was dissolved in 3% DMSO, QNZ-46 from either Tocris or Abcam (UK), methylprednisolone (dissolved in DMSO), minocycline and memantine from Tocris (UK), CP465022 from Viva Bioscience (UK), perampanel (DMSO) from Biorbyt, FluoroMyelin Red (FM-Red) from ThermoFisher Scientific (UK); all other reagents were from Sigma (UK) unless stated otherwise. PLP-GFP and GFAP-GFP mice were maintained as local colonies and tissue-typed using standard protocols.

### Hemi-sectioned brain slices

MCC sections containing the rostral corpus callosum were hemisected in cutting solution and maintained as a matched pair prior to transfer into aCSF for a 60 min recovery period at 37°C. To test the effect of LPS or CPZ, one from each pair of hemi-sections were subsequently exposed to either LPS 1 μg/mL for 60 min or CPZ 1 mM for 120 min. To test the effect of a drug, both hemi-sections were exposed to LPS or CPZ with one of each pair in the presence of a test treatment and the other the drug vehicle. Hemi-section pairs were then fixed in 4% PFA in PBS overnight at 4 °C and stained using FM-Red (1 in 300) for 60 min at room temperature, washed and mounted for imaging. The MCC was imaged on a Leica SPE confocal laser scanning microscope running Leica LAS X Software. Images were subsequently processed and analysed using Leica LAS X and ImageJ. 10x images of the MCC across the hemi-section were collected and tiled, and average fluorescence intensity measured; one pair of hemi-sections from each animal was left unstained and subtracted from the corresponding hemi-sections to correct for the autofluorescence of myelin. 63x images of the corpus callosum were also collected. The protocol is summarized in Ext. data Fig 8.

### CPZ diet

Female C57Bl/6 mice 8 weeks old were fed 0.2 % cuprizone (CPZ) chow (Envigo, UK) *ad libitum* for 1, 2, 4 or 28 days, or were fed CPZ-free matched chow. Mice were weighed and inspected daily with no significant deviation between the control and test groups. Memantine (2mg/kg/day) + perampanel (0.8 mg/kg/day) suspended in 0.5% W/V methyl cellulose was delivered by oral gavage.

### Immunostaining

Tissue was immersed in 4% PFA (Phosphate-buffered saline: PBS pH 7.4) and stored overnight at 4°C prior to sucrose infiltration, embedding (OCT compound, CellPath), and sectioning (20 μm, Leica CM1860 Cryostat). Sections were mounted on Superfrost Plus slides (Thermo Fisher), washed prior to blocking/permeabilization in 5% bovine serum albumin (BSA) and 0.3% Triton X-100 (in Tris-buffered saline: TBS). Anti-LPS: Sections were incubated overnight at 4°C in mouse anti-E. coli LPS (1:200 in 2% BSA and 0.1% Triton X-100 [TBS]) (Abcam; ab35654), washed and incubated in horse anti-mouse biotinylated secondary (1:200 in 2% BSA and 0.1% Triton {TBS]; Vector Laboratories) for 180 min prior to washing and development in Streptavidin-tertiary reagent (1:500 in 2% BSA and 0.1% Triton [TBS]) conjugated to either Cyanine 3 (Cy3) (Merck) or Alexa Fluor 488 (Life technologies) for a further 180 min. Slides were either stained with 4’,6-diamidino-2-phenylindole (DAPI) FluoroPure grade (Invitrogen) and mounted using Fluoromount aqueous mounting medium (Sigma-Aldrich), or underwent further staining procedures. Glia: sections were incubated in either recombinant primary rabbit anti-GFAP antibody (astrocytes, 1:500 in 2% BSA and 0.1% Triton [TBS]; Abcam), or recombinant primary rabbit anti-ionised calcium binding adaptor molecule 1 (Iba1) antibody (microglia, 1:1000 in 2% BSA and 0.1% Triton [TBS]; Abcam) overnight, washed (3x 5 minutes in TBS) and incubated in goat anti-rabbit Alexa Fluor 568 secondary antibody (1:200 in 2% BSA and 0.1% Triton; Invitrogen) overnight at 4°C. After a final wash in TBS, slides were stained with DAPI and mounted as described above. Sections were imaged on a Leica SPE Confocal Laser Scanning microscope running Leica LAS X Software. Images were subsequently processed and analysed using Leica LAS X and ImageJ.

### Image analysis

Cell density was determined blind by manually counting all cells and cells marked by a specific identifier in a set area at 20X magnification using the ImageJ cell counter Plug-in (Kurt de Vos/NIH) to give density as an output per square-millimetre of nerve area. Morphological changes were also analysed, for example as an indicator of the reactivity state of Iba1+ microglia. The characterisation of these states was categorised according to established criteria. Ramified: microglia with distinct ramified processes that extended in an organised manner from the cell body; bushy microglia with truncated processes of varied thicknesses that extended from a swollen cell body; phagocytic (amoeboid) microglia with complete loss of processes and swollen rounded ball-like cell body; motile (rod) microglia with two thin processes that extend from the two poles of the cell body in parallel. Fluorescent intensity and abundance were recorded relative to nerve area (per square-millimetre). Fluorescent intensity was determined as average pixel intensity of a regions of interest; fluorescent abundance was determined as the percentage of nerve area that was positive for immunofluorescent signal. Both were determined using the relevant ImageJ plugins.

### Electrophysiology

MONs were maintained in an oxygenated (1.5 L/min) interface perfusion chamber (Harvard Apparatus Inc.) and continuously superfused (0.6-1 mL/min) with aCSF. They were allowed to recover post-dissection in the chamber for 60 min prior to recording the compound action potential (CAP) using glass electrodes as previously described ^42^. In brief, CAPs were evoked via square-wave constant current pulses (Iso stim A320, WPI), amplified (Cyber Amp 320, Axon Instruments), subtracted from a parallel differential electrode, filtered (low pass: 800–10000 Hz), digitized (1401 mini, Cambridge Electronic Design) and displayed on a PC running Signal software (Cambridge Electronic Design). MONs were crushed at the end of the experiment and the artifact subtracted from the recordings.

### EAE

EAE induction, follow up and motor symptoms score were done as described in detail elsewhere ^52^. Briefly, EAE was induced in C57BL/6 mice by immunization with oligodendrocyte glycoprotein 35–55 (200 μg; Sigma) in incomplete Freund’s adjuvant supplemented with Mycobacterium tuberculosis H37Ra. Pertussis toxin (500 ng; Sigma) was injected on the day of immunization and again 2 days later. Conduction velocity of the corticospinal tract was assessed in anesthetized mice with tribromoethanol (240 mg/kg, i.p.; Sigma) using stimulatory and recording electrodes placed in the primary motor cortex and in the vertebral canal at the L2 level, respectively. After recording, mice were fixed, and spinal cord was processed for histology and immunohistochemistry

### In vivo treatment

CLD-GA treatment; for experimental antagonists, adult C57BL/6 mice were injected i.p. 100 μl of 20 mM β-cyclodextrin in sterile saline + 2 mg/kg QNZ-46 and 1 mg/kg CP465022, or vehicle control without the drugs. QNZ46 and CP465022 were dissolved in DMSO, with a maximal DMSO volume of 4-8 μl (depending on animal weight). Injections were performed blind by animal house staff and mice were left for 240 min on a warming pad. The animals showed no signs of distress or aberrant behaviour.

### Electron Microscopy

Mice were anaesthetized and transcardially perfused for 5 min with cold 4% paraformaldehyde (PFA) in 0.1 M Sørensen’s/PBS solution. The brain was then removed and the MON dissected and immersion fixed in 2.5% glutaraldehyde/0.1 M Sørensen’s overnight and stored in Sørensen’s solution at 4°C. For MCC analysis. Brains were immersion fixed in 2.5% glutaraldehyde/0.1 M Sørensen’s overnight before ∼2 mm blocks were cut from the rostral CC. Spinal cords from EAE mice were isolated and immersion fixed as above. Tissue blocks from the lumbar-sacral region were processed analysed. Isolated MON and MCC subject to the standard LPS or CPZ protocols were immersion fixed in 2.5% glutaraldehyde/0.1 M Sørensen’s overnight and stored in Sørensen’s solution at 4°C. All tissue was osmicated, dehydrated, embedded and thin and ultra-thin (50-70 nm) sections were cut. Ultrathin sections were counterstained with uranyl acetate and lead citrate prior to blind examination using a Jeol JEM 1400 transmission electron microscope. Micrographs were analysed hand via tracing (Image-J, NIH) and the external and internal myelin profile of all X-S axons within a micrograph and converting the area to idealized circles to derive axon diameter and g-ratio. Myelin morphology was blind semi-quantitatively scored for pathology by visual inspection using the following scale: 4= normal myelin morphology, 3= mild myelin pathology, 2= moderate myelin pathology and 1= severe myelin pathology (Ext. data Fig 9).

### Statistics

Data are mean ± SEM, significance determined by t-test or ANOVA with Holm-Šídák post hoc-test as appropriate; Mann-Whitney U test was performed for nonparametric variables. * P≤ 0.05, ** P ≤ 0.01, *** P≤ 0.001. Sample sizes for experimental groups are based on power calculations using established variability. Data from all completed experiments are included and no outliers were excluded. One animal subjected to EAE died and was not counted. All test experiments were intercalated with the relevant controls and where possible trials were conducted blind, including all *in vivo* treatments.

## Acknowledgements

This work was supported by BRACE Dementia Research and the Spanish Ministry of Sciences, (PID2019-109724RB-I00), Centro de Investigación Biomédica en Red, Enfermedades Neurodegenerativas (CIBERNED) (CB06/05/0076) and Basque Government (IT1551-22).

## Extended data

**Extended data Fig 1.**
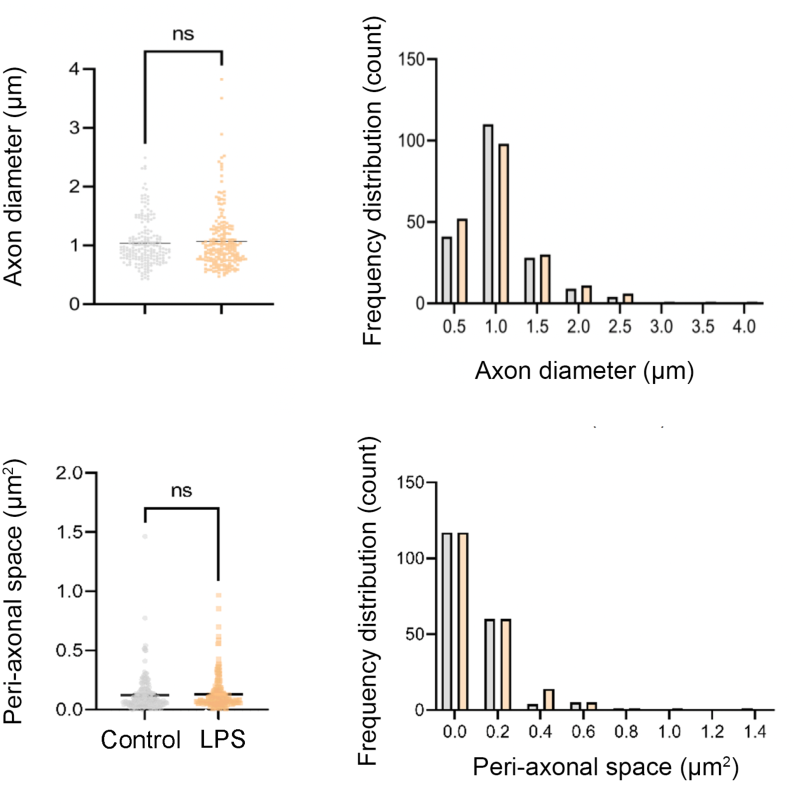
Details of the ultrastructural changes produced by acute 1ug/ml LPS treatment in the MON. Top: LPS (yellow) had no effect upon mean myelinated axon diameter or the size distribution of diameters compared to control (grey). Bottom: LPS had no effect upon the mean area of the peri-axonal space or the area distribution.

**Extended data Fig 2.**
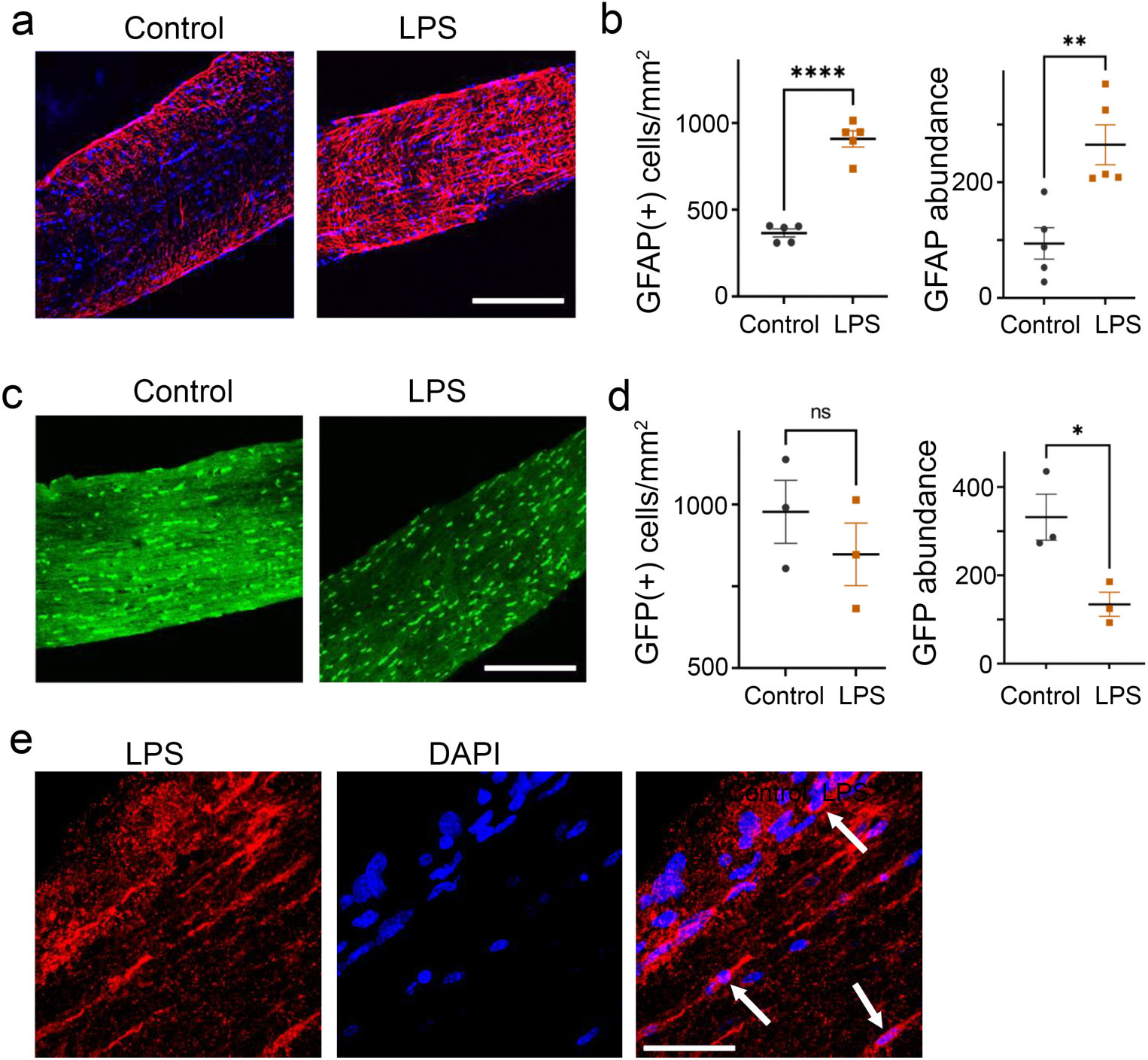
LPS enters the optic nerve and produces reactive astrocytosis. a: Representative confocal images of GFAP immuno-staining (red) of control and LPS treated MON sections (blue: dapi nuclei stain). Scale bar = 50 μm. b: Data summary: there is a significant increase in the number of GFAP(+) astrocytes after the LPS protocol, and in GFAP staining abundance. c: Similar analysis of PLP-GFP MON in the absence of other staining, showing no loss of GFP(+) somata and depletion of staining abundance in the whole sections indicative of process loss. e: Immuno-staining of LPS (red) in MON sections not exposed to other antibodies, showing cell-staining (white arrows). Scale bar = 20 μm.

**Extended data Fig 3.**
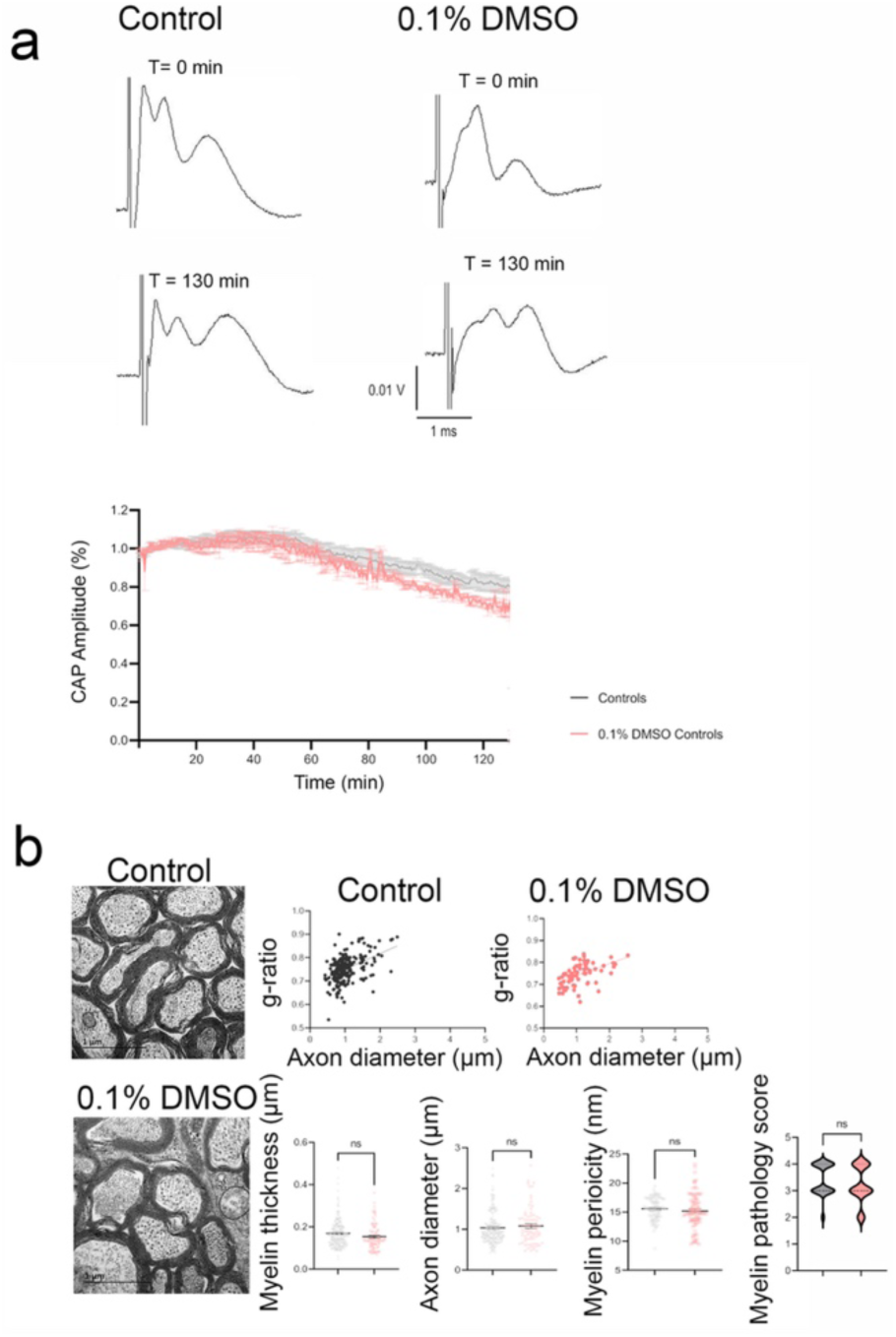
The 0.1 % DMSO vehicle has no significant effect. a: There is no significant difference in the gradual decline in the CAP found during perfusion with aCSF, and during perfusion with aCSF containing 0.1% DMSO (pink). b, left: Representative x-s ultra-micrographs from aCSF perfused (top) and aCSF + 0.1 % DMSO vehicle. Morphometric data is summarised to the right, showing no significant differences in myelin structure between the two conditions.

**Extended data Fig 4.**
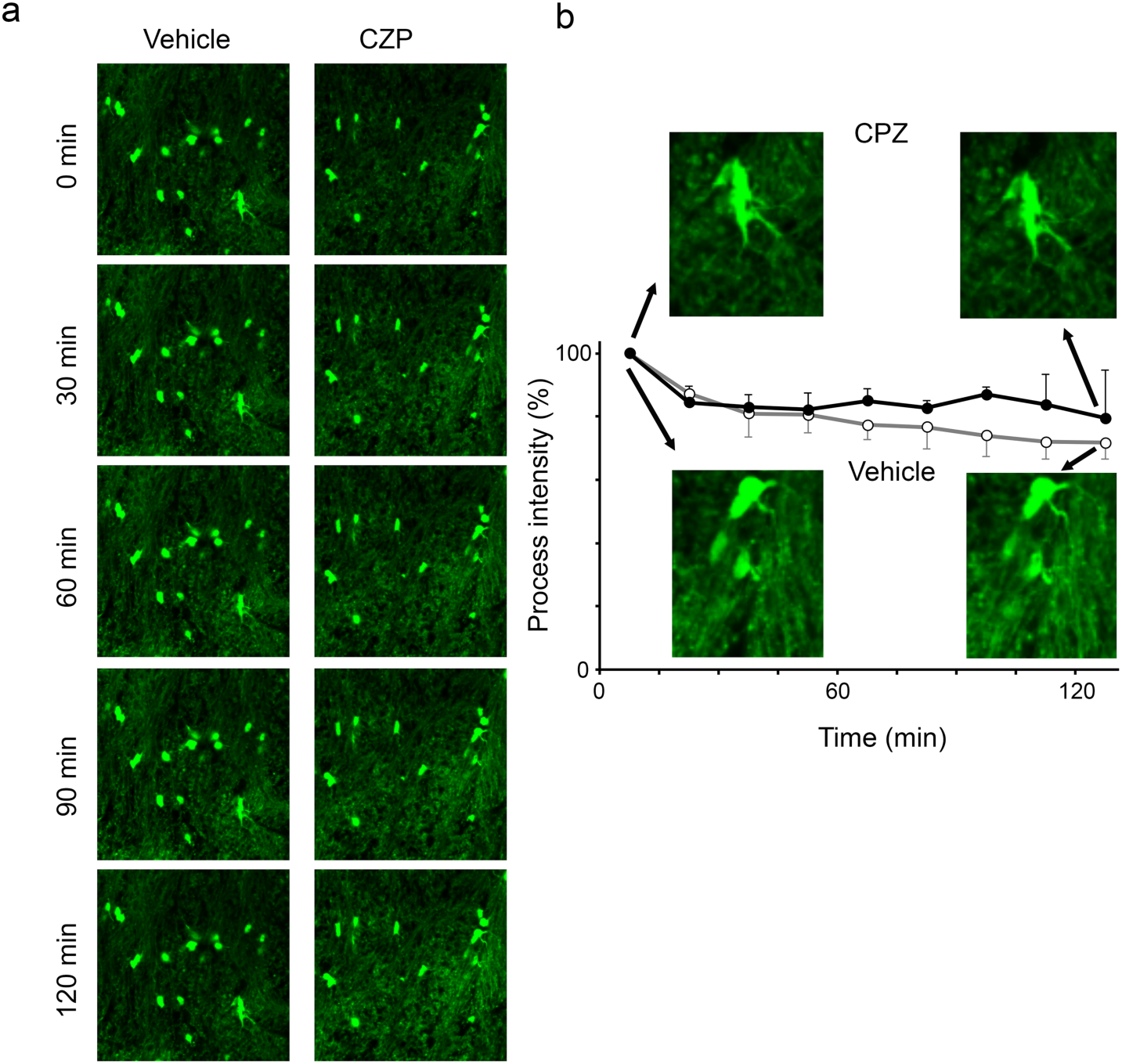
CPZ produces no significant change in oligodendrocyte morphology in the MCC. a: Live imaging of PLP-GFP matched hemi-sections at various time-points (note the preservation of oligodendrocyte number). b: Fluorescent intensity of extra-somata regions shows only a gradual and equivalent loss in vehicle and CPZ-treated hemi-sections. Larger-scale images of individual representative cells are shown in the inserts, illustrating the preservation of cell structure.

**Extended data Fig 5.**
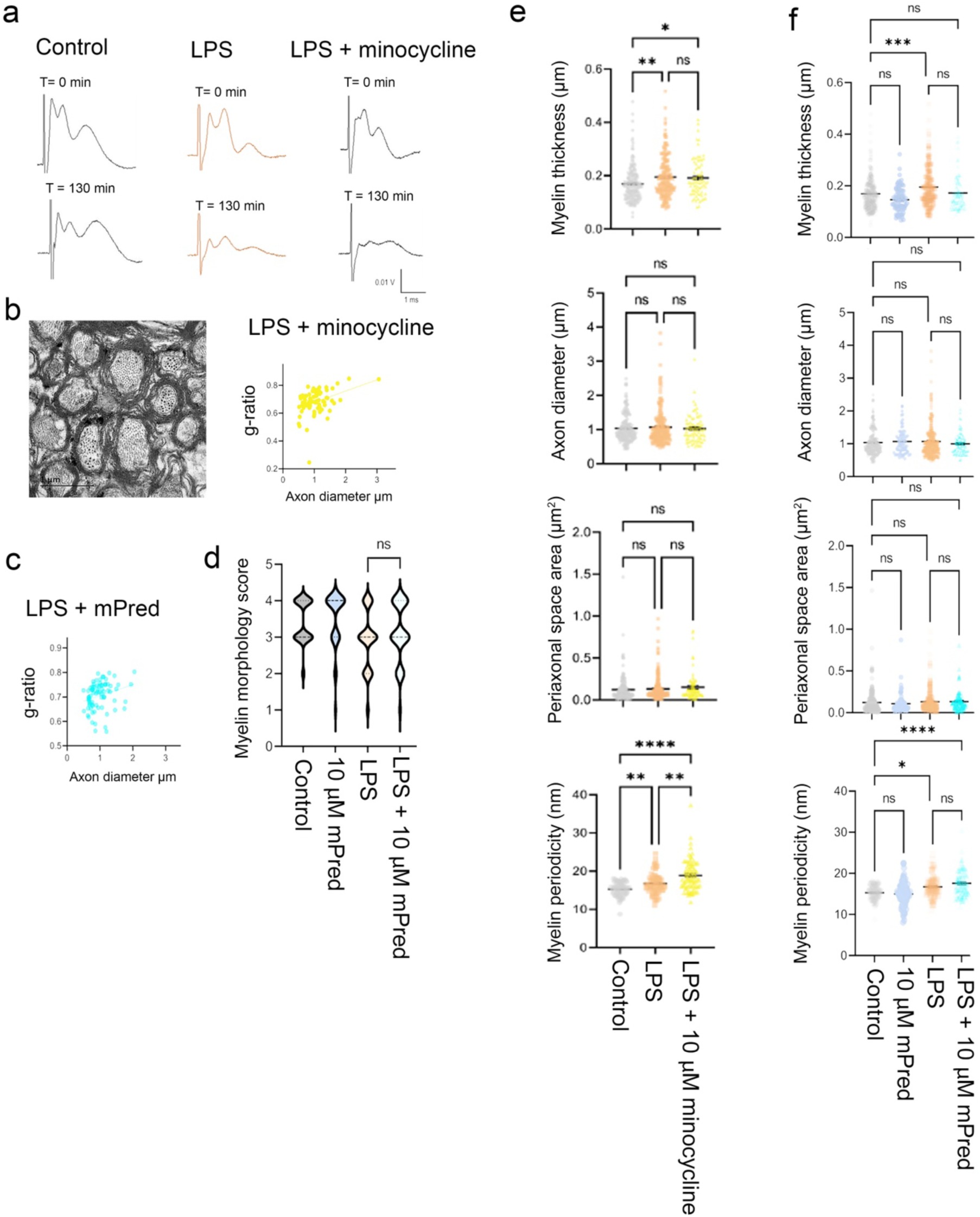
Minocycline and mPred are not protective against LPS-induced myelin injury. a: Example CAPs before and after standard LPS exposure in the presence/absence of 10 μM minocycline. b: (left) Representative electron micrograph from LPS-treated MON in the presence of minocycline. Right: g-ratio vs., axon diameter in LPS + Minocycline treated MONs. c: g-ratio vs., axon diameter in MONs exposed to LPS in the presence of 10 μM mPed. d: Myelin injury score profiles showing no protective effect of 10 μM mPred. e, f: Neither 10 μM minocycline or 10 μM mPred significantly change the effect of LPS treatment upon a range of ultrastructural features in myelinated axons.

**Extended data Fig 6.**
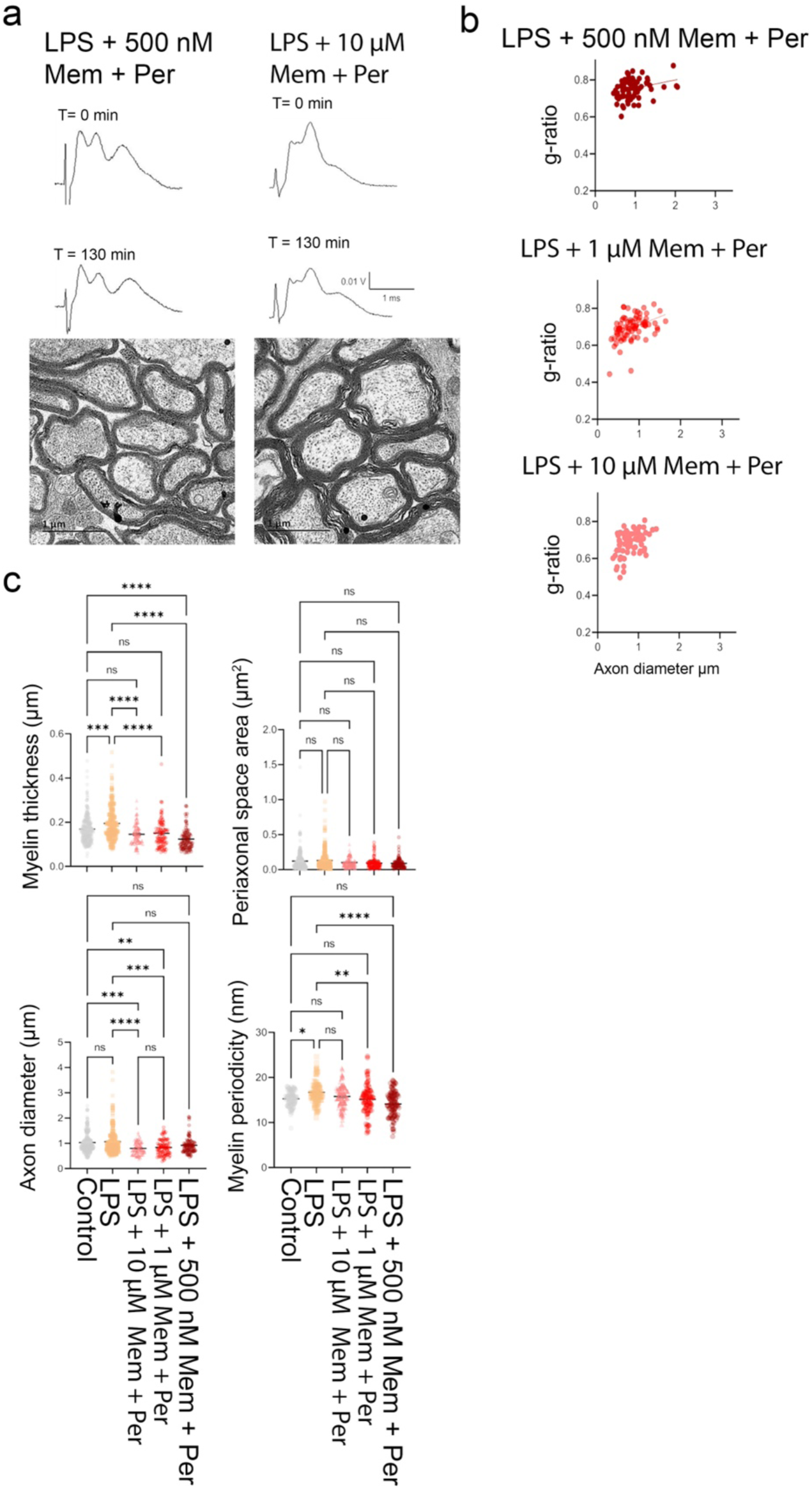
Low dose combined treatment with perampanel and memantine is protective against LPS-induced myelin injury. a, top: Example CAPs before and after LPS exposure in the presence of 500 nM perampanel + 500 nM memantine or 10 μM perampanel + 10 μM memantine. a, bottom: Representative electron micrograph from LPS-treated MON in the presence of 500 nM perampanel + 500 nM memantine or 10 μM perampanel + 10 μM memantine. b: g-Ratio vs., axon diameter for the three combined glutamate antagonist concentrations. c: 500 nM perampanel + 500 nM memantine and 1 μM perampanel + 1 μM memantine significantly protected a range of ultrastructural features in myelinated axons from the effect of LPS treatment.

**Extended data Fig 7.**
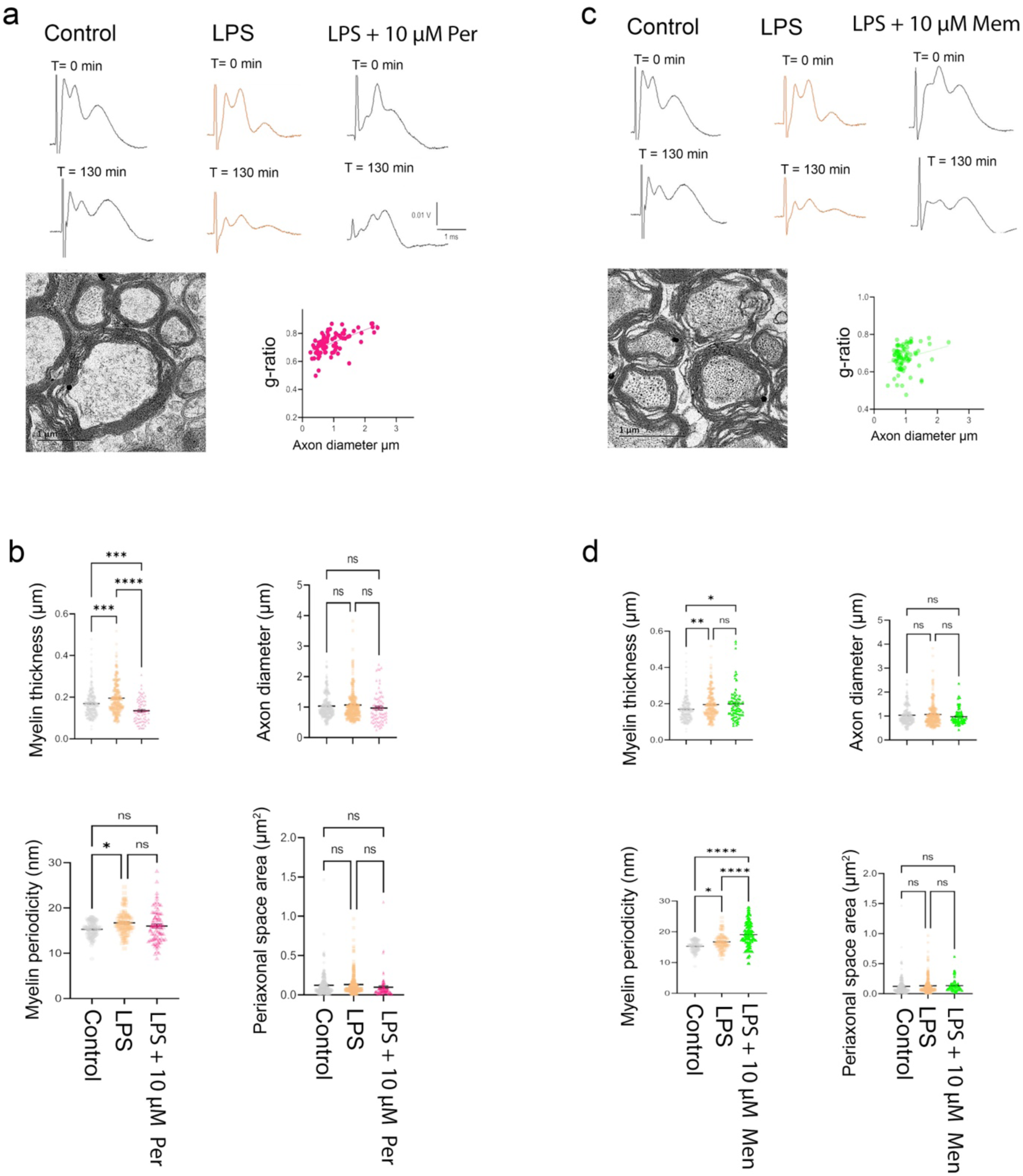
Perampanel and memantine in isolation provide limited or no protection against LPS-induced myelin injury. a, top: Example CAPs before and after standard LPS exposure in the presence of 10 μM perampanel. a, bottom Representative electron micrograph from LPS-treated MON in the presence of 10 μM perampanel and g-ratio vs., axon diameter. b: 10 μM perampanel had limited impact upon the effect of LPS treatment on a range of ultrastructural features in myelinated axons. c, top: Example CAPs before and after standard LPS exposure in the presence of 10 μM memantine. c, bottom: Representative electron micrograph from LPS-treated MON in the presence of 10 μM memantine and g-ratio vs., axon diameter. d: 10 μM memantine did not significantly protect against the effect of LPS treatment upon a range of ultrastructural features in myelinated axons.

**Extended data Fig 8.**
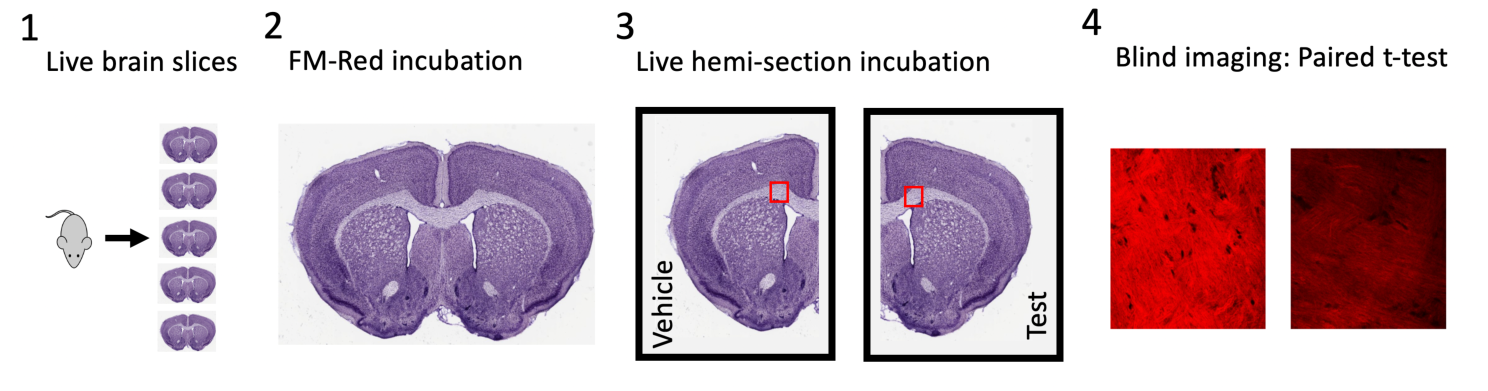
Summary of the acute white matter pathology protocol. 1) Following humane killing and brain dissection, multiple live 120 μm sections are cut via vibratome. 2) Sections are maintained under physiological conditions during a 60 min recovery/FM-Red incubation period. 3) Live sections are hemi-sectioned and maintained under physiology conditions in separate, paired, chambers. One of each of the paired sections may be placed in vehicle media and the other in test media (such as LPS or CPZ), or both may be placed in test media with one of each pair additionally exposed to a test drug. 4). The hemi-sectioned pairs are fixed and the corpus callosum fluorescently imaged under blind conditions, with a paired statistical test used to examine significance.

**Extended data Fig 9.**
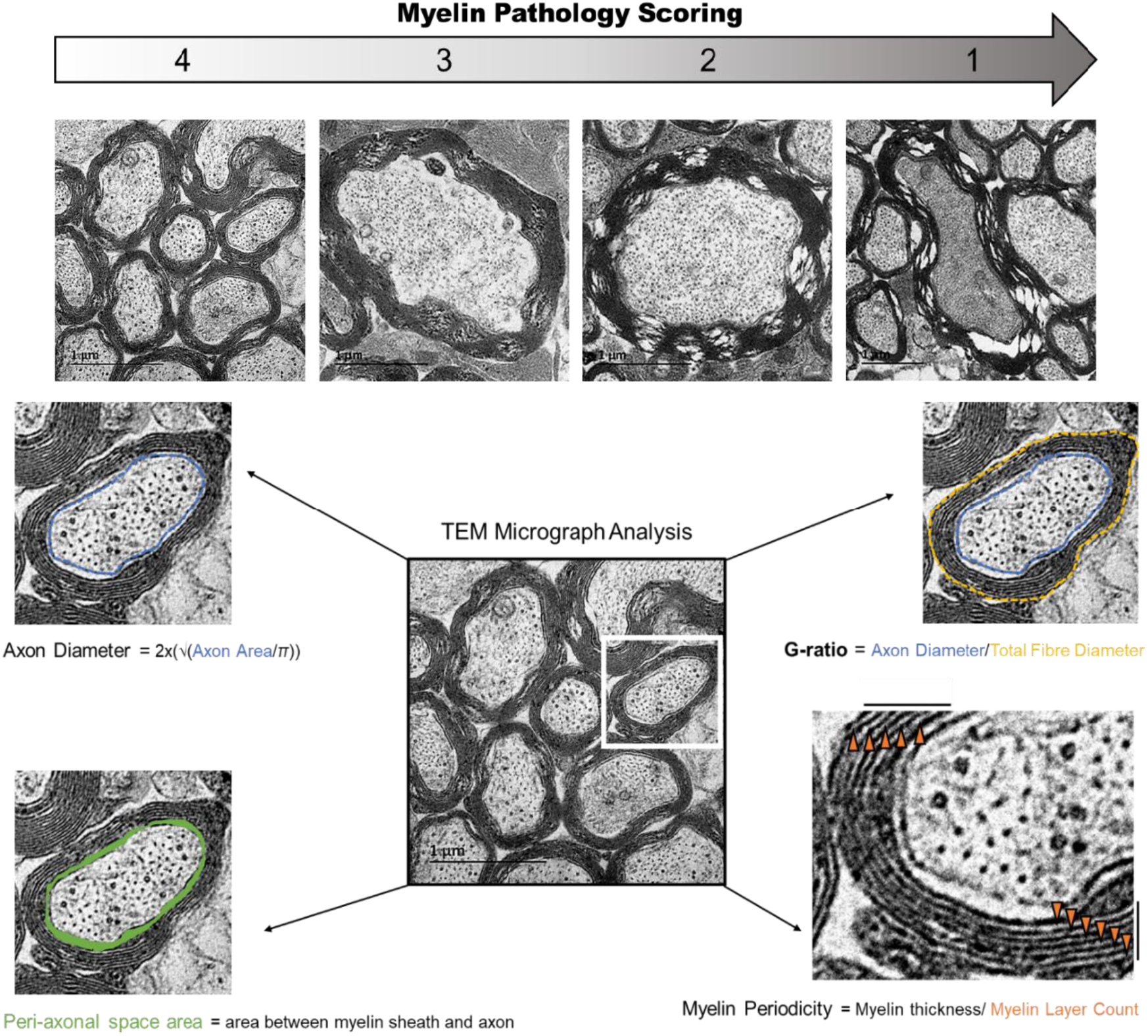
Defining ultra-structural characteristics. Semi-quantitative myelin integrity scoring criteria was defined as a decreasing scale indicating increasing myelin pathology: 4 = no myelin pathology with uniformly compacted myelination, 3 = mild myelin pathology with slight splitting between layers but remaining uniform, 2 = moderate myelin pathology with several points of gross decompaction and loss of uniformity, and 1 = severe myelin pathology where myelin integrity was lost. Key cellular characteristics including axon diameter were defined as idealised axon circles from hand drawn axon area. Peri-axonal space area was calculated by the area enclosed by the myelin sheath and the axon. G-ratio was defined as the ratio of axon diameter to the total fiber diameter. Lastly, myelin periodicity defined as the distance between each myelin layer/wrap was determined by dividing myelin thickness (radius) by the myelin layer count.

